# GAME: Genomic API for Model Evaluation

**DOI:** 10.1101/2025.07.04.663250

**Authors:** Ishika Luthra, Satyam Priyadarshi, Rui Guo, Lukas Mahieu, Niklas Kempynck, Damion Dooley, Dmitry Penzar, Ilya Vorontsov, Yilun Sheng, Xinming Tu, Adam Klie, Shiron Drusinsky, Alexander Floren, Ethan Armand, Kaur Alasoo, Georg Seelig, Ryan Tewhey, Peter Koo, Vikram Agarwal, Sager Gosai, Luca Pinello, Michael A. White, Avantika Lal, Julia Zeitlinger, Katherine S. Pollard, Maxwell Libbrecht, Hannah Carter, Sara Mostafavi, Ivan Kulakovskiy, Will Hsiao, Stein Aerts, Jian Zhou, Carl G. de Boer

## Abstract

The rapid expansion of genomics datasets and the application of machine learning has produced sequence-to-activity genomics models with ever-expanding capabilities. However, benchmarking these models on practical applications has been challenging because individual projects evaluate their models in ad hoc ways, and there is substantial heterogeneity of both model architectures and benchmarking tasks. To address this challenge, we have created GAME, a system for large-scale, community-led standardized model benchmarking on user-defined evaluation tasks. We borrow concepts from the Application Programming Interface (API) paradigm to allow for seamless communication between pre-trained models and benchmarking tasks, ensuring consistent evaluation protocols. Because all models and benchmarks are inherently compatible in this framework, the continual addition of new models and new benchmarks is easy. We also developed a Matcher module powered by a large language model (LLM) to automate ambiguous task alignment between benchmarks and models. Containerization of these modules enhances reproducibility and facilitates the deployment of models and benchmarks across computing platforms. By focusing on predicting underlying biochemical phenomena (e.g. gene expression, open chromatin, DNA binding), we ensure that tasks remain technology-independent. We provide examples of benchmarks and models implementing this framework, and anticipate that the community will contribute their own, leading to an ever-expanding and evolving set of models and evaluation tasks. This resource will accelerate genomics research by illuminating the best models for a given task, motivating novel functional genomic benchmarks, and providing a more nuanced understanding of model abilities.

## Main

While sequence-based genomics models are widely used in the field, their progress has proven difficult to benchmark due to substantial heterogeneity in model architectures, training datasets, and ad hoc model evaluations. Presently, model evaluations are siloed within publications, resulting in duplicated efforts; simultaneous differences in model architectures, training data, and evaluation tasks often obscure where any innovation lies. In general, functional genomics assays measure the genome’s biochemical activity indirectly. Consequently, some performance gains are likely attributable to the models capturing assay-specific biases better, rather than capturing the underlying biological phenomenon. Further, it is commonplace to use genomics models to predict activities that are related to but distinct from the tasks they were designed to predict (e.g. using CAGE predictions for mRNA expression even though they capture different RNA subsets)^1^. Finally, even if a benchmark is superficially the same, there are often many choices (degrees of freedom) in implementation that may impact the performance measure. Consequently, the overall performance of a new genomics model compared to its predecessors and contemporaries often remains opaque. Recent work showed that proper and consistent benchmarking can reveal model limitations^2–6^. For instance, Sasse et al (2023) showed that, while models are able to predict the magnitude of variant effects, they struggle with predicting the direction of the change in expression^3^. As the number of potential benchmarking datasets continues to grow, facilitating their uniform application across models becomes increasingly important.

In other fields, there are well-established, high-quality datasets that can be used to benchmark progress because it is generally agreed that they capture the task well. For instance, Critical Assessment of Structure Prediction (CASP)^7^ is used to benchmark protein folding models, while the CIFAR datasets are standard benchmarks for image classification^8^. Similarly, there have been efforts to curate diverse sequence datasets into a standard format to facilitate model training and evaluation^5,9^. While challenges and competitions like DREAM^10,11^ and Kaggle^12^ are excellent at determining the optimal model design choices for specific datasets, they are, by their nature, restricted to specific tasks. However, there are many distinct but intertwined activities encoded by the genome (e.g. RNA expression, open chromatin, transcription factor binding, histone marks, 3D conformation) that differ across species, cell types, and conditions, making it necessary for sequence-based genomics models to be benchmarked on tasks that match the model’s scope and purpose. Further, users may want to identify the best model for their specific task of interest.

To enable systematic benchmarks of various models by various evaluation tasks, we created GAME (**G**enomics **A**PI for **M**odel **E**valuation), a framework for large-scale community-led standardized model benchmarking. GAME is designed to be flexible and user-friendly, where new and existing models can be systematically and uniformly queried with an ever-expanding variety of tasks. To facilitate this uniform benchmarking of models across tasks, we leveraged the standardized communication protocols used by APIs^13,14^. APIs enable diverse client programs to interact with server programs, each without knowing any of the details of the other’s implementation, facilitated by the common language defined by the API protocol. While Avsec et al. (2019) created a repository, Kipoi, for genomics-based ML models and also used APIs to access them, it did not include benchmark modules, and it required the user to have intimate knowledge about exactly what each model was predicting, making scalable, uniform model benchmarking a substantial challenge^15^. GAME allows for seamless communication between pre-trained models and functional genomics datasets on which those models can be benchmarked. GAME was designed with extensive community feedback throughout its development, resulting in API specifications that meet the needs of model and dataset experts.

To maximize GAME’s utility and adaptability, we established several forward-looking design principles that emphasize creating benchmarks that are as objective and biology-focused as possible. Firstly, we focused on the biological signals of interest, not the assay used to quantify them. Accordingly, assay types are not provided to the models, which could use this information to improve performance but in biologically meaningless ways (e.g. adding assay-specific biases). For example, DNase I-seq and ATAC-seq both measure chromatin accessibility in different ways and have different biases. While predicting the assay-specific biases would increase performance, it provides no insight into the underlying biology. This has the secondary benefit of being robust to the development of new assays, which are likely to measure the same underlying biological phenomena in different ways. For instance, the recently-developed Fiber-seq also measures chromatin accessibility, but the nature of the data and biases differ substantially from DNase I-seq and ATAC-seq^16^. Accordingly, we restrict the predictions to DNA binding (which can be for TFs, methylation, or histone marks), chromatin accessibility, 3D-chromatin conformation, and expression (mRNA, Pol I, Pol II, Pol III). Secondly, we aimed to provide the utmost separation between the prediction and evaluation of these models by providing models only as much information as needed to perform their tasks and no more to prevent overfitting to the benchmarks. For instance, predicting a specific biological replicate could increase performance (e.g. if the model saw the replicate during training), but would be misleading because it would not generalize to the samples we actually care about. Thirdly, we aimed to make this framework sustainable in the long term by distributing responsibilities to those making models and benchmark datasets, and having minimal centralized maintenance. New models/datasets need only to implement the API to seamlessly communicate with all others that already exist in the framework. While this adds some initial overhead, the existing models and benchmarks in GAME will mean that implementing the GAME API will become the most efficient way to complete evaluations.

In GAME, models are encapsulated in Predictors and benchmarks are encapsulated in Evaluators, which communicate via the API protocol. The Evaluator module is a software client that is responsible for evaluating models on a specific benchmark, while the Predictor module is a server that listens for incoming requests and responds with the model’s predictions for the requested tasks. We also have included a Matcher module which uses standardized queries to a local LLM^17^. Matcher is responsible for mapping requested tasks with tasks the model knows how to do (e.g. species, cell types, and molecules). For example, the Matcher must map the cell types requested by the Evaluator to the closest matched cell types that the Predictor is able to predict. The Matcher is implemented as another server for which the Predictor serves as the client. A GAME evaluation occurs in 7 steps (**Fig. 1**). (1) The Evaluator requests predictions from a Predictor. (2) The Predictor parses the request and, if needed, consults the Matcher to determine how it can best serve up the tasks requested. (3) The Matcher finds the mapping between the requested tasks and tasks the predictor can do. (4) The Matcher sends this information back to the Predictor. (5) The Predictor processes the request, by sending it to the model and reformatting the model’s predictions in the required standard format. (6) The predictions are returned to the Evaluator. (7) Finally, the Evaluator calculates the model’s performance. The Evaluator builders are responsible for determining what information the Predictors require for their predictions and what the best evaluation metrics are for their benchmarks. The Predictor builders are responsible for taking a request for predictions on a set of DNA sequences and determining how it can best fulfill the request, given what the model can predict (or decline to predict if it cannot do so). The Evaluators, Predictors, and Matcher are containerized using Apptainer to facilitate sustainable model evaluation across computing environments, improving our ability to compare models^18^. Inside the containers, the modules can be written in any language. However, to facilitate the adoption of this approach, we provide examples and templates for functions implemented in Python with seamless GPU support that users can modify to fit their needs.

**Figure 1.**
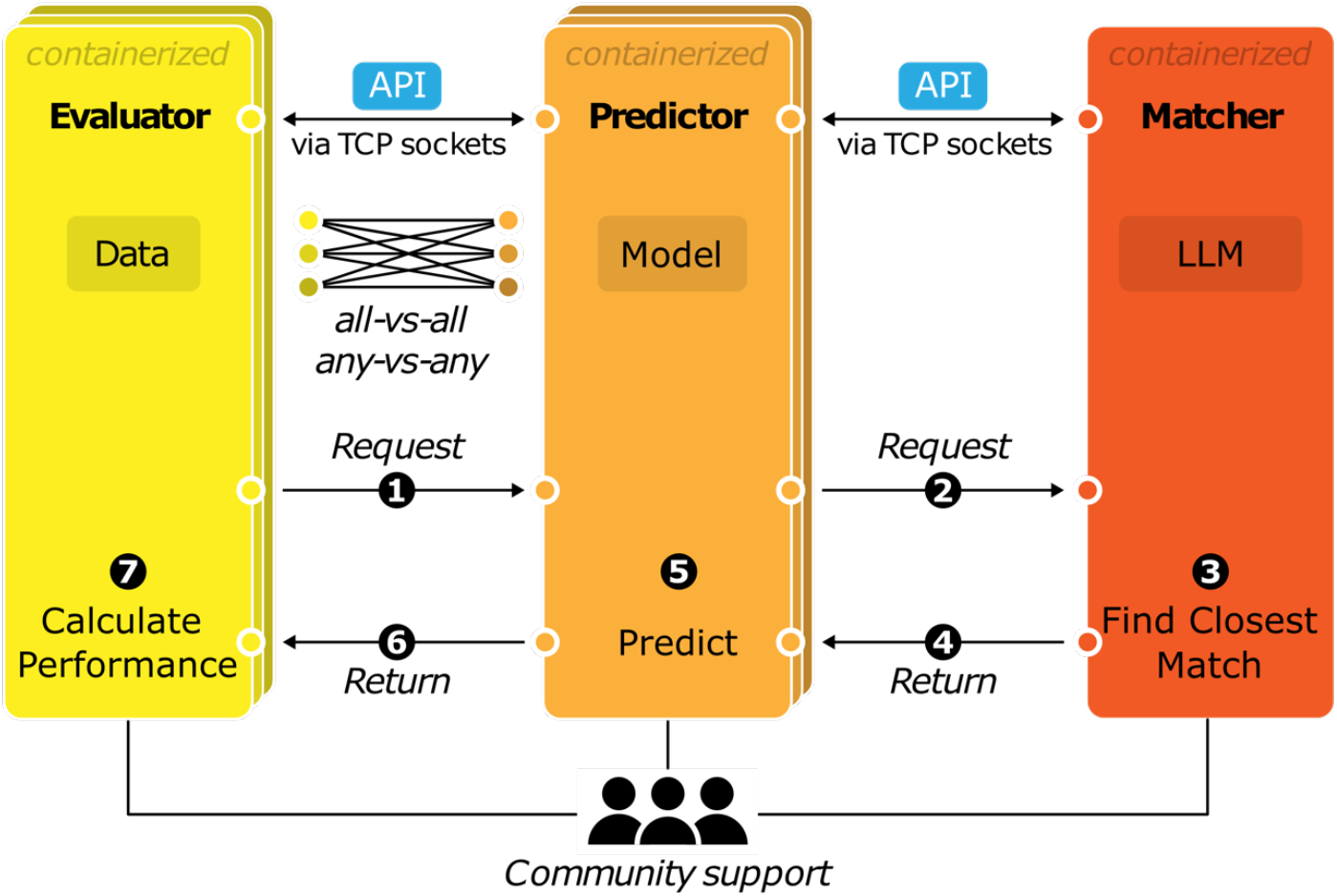
GAME framework. GAME includes three modules: The Evaluator, containing a benchmark dataset; the Predictor, encompassing a sequence-to-activity model; and the Matcher, capturing relationships between tasks. All GAME modules are inherently interoperable by communicating in the GAME API protocol over TCP. For each benchmark, the Evaluator requests a prediction from the Predictor, which consults the Matcher to determine the closest task the Predictor can complete. Once the Matcher returns the best match, the Predictor will complete its prediction and return it to the Evaluator, which will evaluate performance. Members of the genomics community will contribute modules to enable continual evaluation of more models across more benchmarks.

Including a separate Matcher module provides several advantages. The cell types requested by the Evaluator are likely imperfect matches to the cell types the Predictor can predict (e.g. cardiomyocyte vs. heart or HEK293FT vs. HEK293). Similarly, the Predictor may not include the requested species and so would need to know which of the species it includes is closest. The Matcher resolves such discrepancies by automating the alignment of tasks. This ensures that performance measures are not influenced by the Predictor builder’s understanding of cellular, molecular, or species relationships. Further, as our understanding of species and cell types evolves, the Matcher module can be upgraded while remaining compatible with existing GAME modules.

We applied GAME to several models and two representative benchmarking tasks – gene expression and chromatin confirmation (**Fig. 2**). For gene expression, we implemented three Evaluators, two using MPRA datasets (both genomic and synthetic)^19,20^, and one of measured effects of synthetic variants^21^, each of which can output distinct evaluation metrics. For example, the Agarwal et al. MPRA Evaluator evaluates models on their ability to predict gene expression separately for each cell type, and also on how well models predict cell type-specific differences in expression (**Fig. 2a**)^19^. The Gosai et al. MPRA Evaluator data includes synthetic (non-genomics) MPRA probes, and so is an ideal benchmark for genome-trained models. Similarly, the Martyn data represents the expression effects of synthetic genetic variation introduced by genome editing, and so is a good task benchmarking models on variant effects with minimal risk of train-test leakage. We created Predictors for Borzoi, Enformer, and DREAM-RNN, and evaluated each on each expression Evaluator (**Fig. 2a**). Borzoi and Enformer are both large sequence window genome-trained state-of-the-art sequence-to-function models and have been used in numerous benchmarking papers^22,23^. The DREAM-RNN Predictor is a state-of-the-art short-sequence sequence-to-function model^24^, in this case trained on Agarwal K562 MPRA data^19^. As a more complex example we also created a 3D chromatin confirmation (**Fig. 2b**) Evaluator and Predictor based on

**Figure 2.**
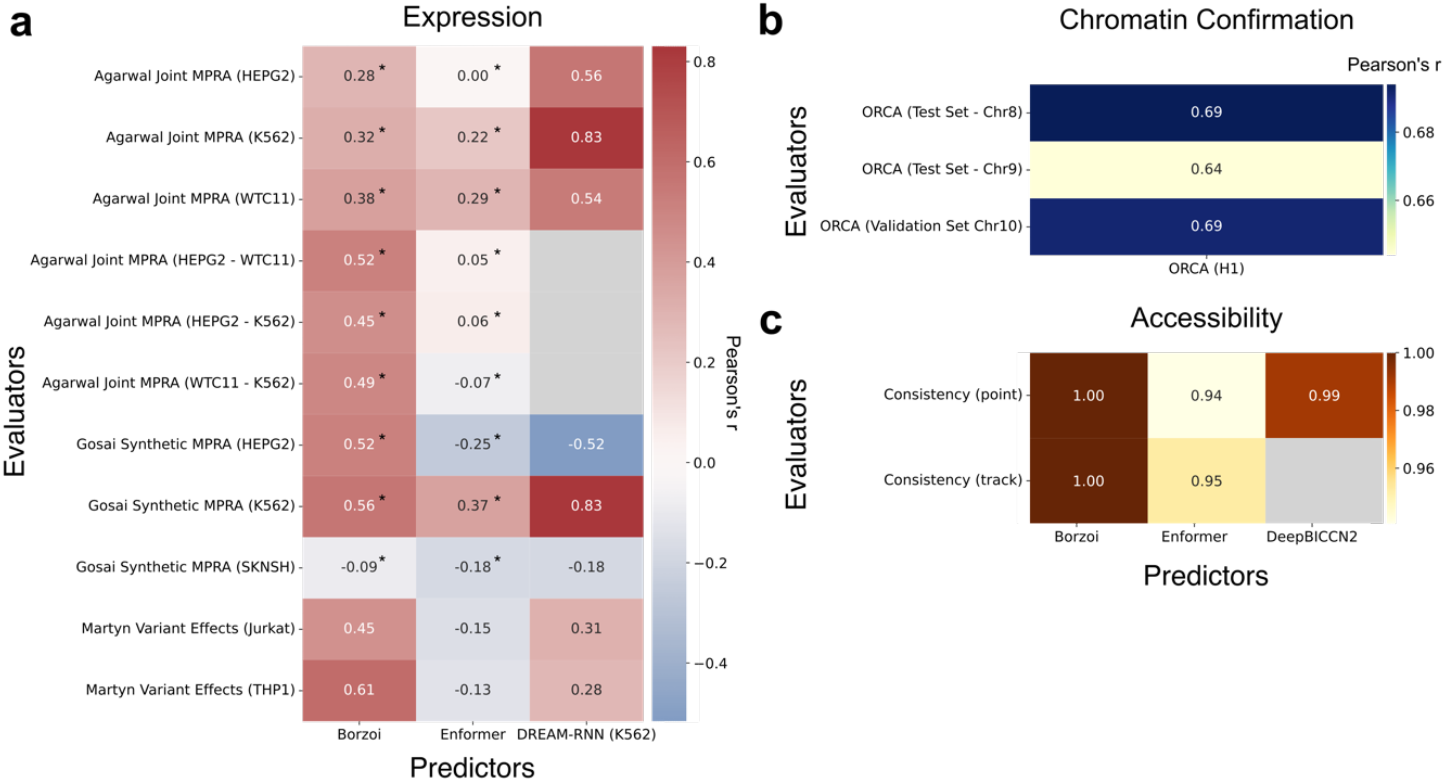
Sample benchmarking done with GAME. **a**, Expression evaluation tasks. Models (*x* axis) were evaluated for their correlation to measured expression levels (colours) across a variety of tasks (*y* axis). **b**, Chromatin conformation tasks. Correlation of Orca predictions vs measured chromatin contact frequencies (colours) for two Orca test-set chromosomes and one validation-set chromosome (*y* axis). **c**, Consistency evaluation for accessibility. Point and track based accessibility consistency evaluators (*y* axis) were used to evaluate the correlation between predictions for forward and reverse complement sequences (colours) in models of DNA accessibility (*x* axis). *Correlation values from down sampled datasets. Grey cells mark values that were not calculated due models that could not complete the Evaluator’s requested task.

Orca^25^. Finally, we developed Consistency Evaluators that can be used to assess models that predict DNA accessibility, confirming correlated predictions between forward and reverse complement sequences (**Fig 3b**) for Borzoi, Enformer, and the recently-described ATAC-seq model DeepBICCN2^26^

GAME offers many advantages that will streamline benchmarking and model use more generally. The common protocol enables any model to communicate with any benchmark (although not all combinations may make sense), ensuring that when one creates a new GAME module, it is immediately compatible with all the existing modules. Existing Predictors and Evaluators remain frozen so that future benchmarks done with them are identical; future evaluations of models on these benchmarks will be directly comparable to the results shown in **Fig. 2**. The anonymity of API requests means that all models will be evaluated equally.

The modules communicate via TCP/IP sockets and so can be run locally or remotely, potentially preserving privacy, for example, in cases where models have proprietary weights. Because GAME is decentralized and community-driven, it will be much more scalable and easily maintained. Contributors can easily add their own benchmarks and models, and can maintain them as long as they remain relevant. We have already implemented 5 models and 6 benchmarks in GAME.

As GAME gains momentum, the fastest and easiest way of performing a new benchmark will be to implement GAME, further incentivizing this community effort. We anticipate that GAME will drive genomics progress in several ways. GAME allows one to easily compare all models across all benchmarks, which will reveal model limitations and low-quality benchmarks. For instance, a model that was overfitted to a current benchmark would be revealed as it fails to generalize to the benchmarks that arise after its release. Such a framework also facilitates continual evaluation of state-of-the-art models on a diverse set of tasks, enabling researchers to quickly identify the models that are most well-suited to their specific tasks. For instance, if one is interested in predicting the effects of genetic variants in heart, one can find the closest benchmark to determine which model is best for the task. To showcase GAME in a reasonable time frame with reduced computational costs, we down-sampled certain Evaluator datasets (**Fig. 2**). However, Evaluators and Predictors can split up the dataset, enabling parallelization and better utilization of available resources. While we anticipate that Matcher will be useful outside of GAME, determining the best way to match tasks turned out to be a substantial challenge. Nevertheless, the Matcher can be continually improved due to its modular nature. Finally, having all models share a common language means that other software can use GAME to facilitate model reuse and swapping between models more easily. Its modular design will keep GAME in play, evolving alongside the ever-changing field of genomics.

## Methods

### Information passing

The client and server code of an API can be written in any programming language^27^. However, due to the prevalence of Python programming in genomics and model development^28,29^, the GAME API and corresponding error-checking functions are written in Python. Python sockets can connect servers to clients to communicate with each other over local or remote networks^30^. Python sockets use Transmission Control Protocol (TCP), which provides benefits such as detecting lost information when sending information over a network and controlling the order of data when transmitted^31^.

The Predictor server will bind to a user-specified “HOST,” an IP address or hostname, and “PORT,” a TCP port. Once the server binds, it will wait and listen for incoming connections. The Evaluator client must connect to the same HOST and PORT pair to send a request to the server. The data that is transmitted via the TCP socket must be converted to bytes before it is sent. The Evaluator and Predictor can both be on a local computer, HPC platform, or even in remote configurations, ensuring flexibility in how they communicate. This capability is also beneficial for Predictor developers working with proprietary or closed-source models, allowing them to integrate their work into the GAME framework. Those models can be embedded into the Predictor codebase and hosted as a server on their own network, maintaining full control and privacy over their model weights.

### Communication protocol

In GAME, the communication protocol distinguishes between the initial format of user-provided data and the standardized formats used for data streaming between GAME modules. While users can store benchmark data in various initial file formats (e.g. .json, .npy, .txt, .xlsx), these inputs must first be converted to the correct format before transmission. Most typically, this is implemented as a helper function that converts the input file into JavaScript Object Notation (JSON)-like dictionary structure. It is essential that this structure adheres to the predefined JSON schema of GAME to ensure data compatibility and proper interpretation by subsequent modules. Data in the JSON object is stored as key: value pairs^32^.

Once the data is in this standardized dictionary representation, it is ready for transmission. By default, GAME utilizes JSON for this inter-module data exchange, due to its popularity, speed in generation and parsing, and support via Python’s standard library^33^. We also allow information to be transmitted using MessagePack (MsgPack), a binary serialization format, which helps increase speed when returning large arrays of predictions^34,35^, provided both the Evaluator and Predictor modules are configured to handle it. The Evaluator, therefore, serializes the prepared dictionary structure into either a JSON or a MsgPack stream before sending the required information to the Predictor. If the task requested is not an exact match to one of the Predictor’s model outputs, the Matcher module can be utilized to align the requested cell types, species, and molecules with those supported by the Predictor. Once the prediction is complete, the predictions are reformatted to the negotiated return format and returned to the Evaluator. JSON and MSGPACK files are easily loaded into Python and accessed as dictionaries, making them straightforward to work with^36^. For implementation guidance, sample files and code can be found on GitHub.

### Module responsibilities

The Evaluator and Predictor modules are designed to have specific responsibilities that follow the core principles of the API, such as minimizing information sent to a Predictor, allowing dataset creators the power to evaluate the tasks they are interested in, and allowing model builders the freedom to parse sequences to comply with model specifications as they see fit. The main responsibilities of Evaluators are to choose what predictions they would like to obtain from the Predictor and what information the Predictor needs to fulfil that request, including: whether the Predictor should return points (a single value) or tracks (values across a sequence), whether there are constant flanking sequences that could provide relevant context (e.g. the vector backbone of an MPRA vector), where in the sequence window the predictions should be provided (e.g. the location of a promoter), how to use the expression predictions based on sense or antisense strand. This information is encoded in the request that is sent to the Predictor. Predictors, conversely, are responsible for parsing the sequences to meet model input size requirements (pad, trim, or add other flanks to the sequences), aggregate across tracks (e.g. replicates of an experiment) or bins (e.g. if the prediction window overlaps multiple bins), deciding which cell type/species/molecule to use (with help from the Matcher, if desired), and returning the predictions in the standard format. Once the predictions are returned, it is the Evaluator’s responsibility to calculate the appropriate metrics for each benchmarking task and output these results to a file, using a standard reporting format. Additional details of the API specification and examples are available on GitHub.

### Containerization

Containers provide a secure and robust way to run code across any platform, removing troublesome dependency issues. Since most ML models are trained and evaluated on high-performance computing (HPC) platforms, we used Apptainer for GAME module containers. Unlike Docker, Apptainer is specifically designed for HPC environments and does not require root access, a critical advantage since most HPC systems restrict root privileges for security reasons^18^. By containerizing each Evaluator and Predictor, the code within each container is isolated, enabling each to use its own version of any language without issue. Additionally, in contrast to Docker, Apptainer containers have no network isolation, meaning that they can communicate with each other using IP addresses and ports without having to create a network for them to communicate over.

### Maintenance and Community Contributions

The long-term maintenance of this framework is designed to be simple and scalable. We have made community-maintainable lists of GAME modules available on GitHub, which then needs only to be updated with the locations of new modules. For instance, someone could create a new model, package it into a Predictor, and then issue a pull request to update the list of Predictors with their own. Subsequently, users can pull the latest lists of Predictors and test them on the latest list of Evaluators. The module list includes their names, a description of their function, and links to the locations where containers are stored. Zenodo provides an ideal location for housing Evaluator and Predictor containers because it is durable and has the additional benefit of scanning uploads for viruses to ensure the safety of containers for users to pull^37,38^.

### Automated and externalized cell type, molecule, and species matching

GAME includes a module called “Matcher” that automatically maps the Evaluator’s requested species, measured molecule, and cell type with what a Predictor can provide. The Matcher is designed to perform this task by interpreting the relationship between terms through lexical, syntactic, and sematic matching. Lexical matching handles cases of direct string correspondence, such as finding the exact token ‘A549’ within a more descriptive choice like ‘lung adenocarcinoma cell line: A549’. Syntactic matching addresses structural variations and common abbreviations, such as ‘hek-293’ or ‘SKNSH’ to ‘HEK293’ or ‘SK-N-SH’, respectively. Semantic matching uses biological knowledge to connect different terms that refer to the same entity, such as mapping the description ‘chronic myelogenous leukemia cell line’ to a canonical cell line name, ‘K562’.

The matcher uses a local large language model – Gemma 3, which is a collection of very lightweight and efficient models that can run effectively on a single GPU or TPU^17^. Running Gemma 3 locally means that the model operates directly on the hardware of the system it is deployed in, ensuring data privacy, reducing reliance on external cloud APIs, and lowering operational costs, which are crucial for sensitive or high-throughput workflows.

The Matcher module utilizes Ollama to run and manage Gemma 3 locally^39^. Ollama is an open-source platform that simplifies hosting large-language models (LLMs) on local hardware by packaging models and providing tools for their management. Ollama helps abstract away many underlying complexities of model backends using inference engines and can automatically leverage available GPU resources for accelerated performance. This ease of setup and serving is key to the containerized and reproducible design of the Matcher module^40,41^. Additionally, the availability of an official Docker image for Ollama further streamlines the creation of portable and reproducible runtime environments. When a model is run, Ollama exposes a local API endpoint (on port 11434). This allows applications, like the Predictors in GAME, to send requests to the LLM and receive responses.

To enable the Matcher module Python environment to interact with the Gemma 3 model, the LangChain framework is utilized^42^. LangChain is an open-source library designed for developing applications that incorporate LLMs^42,43^. It provides a collection of tools that assist in managing prompts, interacting with LLM APIs (including local instances like those served by Ollama), and constructing more extensive LLM driven pipelines. Within the Matcher module, the `langchain-ollama` package provides specific client classes, like `OllamaLLM`, which facilitate communications with the local Ollama API. This integration enables the Matcher to send structured input queries – comprising the fuzzy input term (Evaluator request) and the list of choices (what the Predictor can provide predictions on) – to the Gemma 3 model through the Ollama server and then to receive the best identified match. To create structured queries consistently, the `ChatPrompTemplate` class is employed from LangChain, which supports the goal of reproducible matching. By automating this process, we reduce potential biases due to memorization of cell-type relationships. This also circumvents the process of manual selection or literature review to select closest cell types. Matcher can also be used to align species or tasks (e.g. ‘NF-kB’ to ‘NFKB1’).

A key challenge in this process is that a Predictor may have hundreds of available choices, a list too long to fit within the context window of the LLM^44,45^. To overcome this, the Matcher implements a chunking strategy. The complete list of choices is first divided into smaller, manageable chunks. The LLM then performs a ‘tournament-style’ elimination, finding the best possible match, the ‘champion’, within each individual chunk. Finally, a championship round is held where the LLM evaluates only the list of champions to determine the single best overall match. This method makes the matching process scalable to arbitrarily long lists of choices.

## Data and Code availability

API specifications, code and additional documentation can be found on GitHub (https://github.com/de-Boer-Lab/Genomic-API-for-Model-Evaluation/tree/main). List of current Modules can be found here: https://github.com/de-Boer-Lab/GAME_modules

## Acknowledgements

We would like to thank J. Gagneur, T. Mauermeier, J. Shendure, C. Qiu, D. Calderon, A. Gao, P. Fradkin, B.J Frey, G. Eraslan, L. Gunsalus, S. Nair, D. Kelley, R. Das, S. Kundu, I. Raine, V. Hecht, A. Kundaje, Z. Avsec, J. Engreitz, A. Gschwind, M. Montgomery, M. Weilert, C. McAnany for helpful feedback and discussion during the API design and implementation process. We would also like to thank M. Krzywinski for design help with Figure 1. M.A.W was supported by National Institutes of Health (USA) R01GM121755. J.Z, S.D and A.F were supported by Additional Ventures, Keck Foundation, Biswas Family Foundation, Gladstone Institutes. J.Z was supported by Cancer Prevention and Research Institute of Texas grant RR190071 and National Institutes of Health (NIH) grants DP2GM146336.

## Author Contributions

I.L and C.G.D conceived and designed GAME. I.L and S.P set up GAME code base and documentation. R.G set up codebase for Orca Evaluator and Predictor. L.M and N.K contributed DeepBICCN2 Predictor. M.M provided data for variant effects Evaluator. I.L, S.P, and C.G.D wrote the paper. K.P, H.C, A.K, J.Z, L.P, M.A.W, A.L, I.K, I.K, D.P, and K.P provided feedback on the manuscript. D.D and W.H provided feedback for Matcher and helped create early implementations of the Matcher module. All other authors were involved in either the design process for the API specifications, feedback and/or testing API tutorials and setup.

